# Developmental plasticity of texture discrimination following early vision loss in the marsupial *Monodelphis domestica*

**DOI:** 10.1101/2020.09.05.284554

**Authors:** Deepa L. Ramamurthy, Heather K. Dodson, Leah A. Krubitzer

## Abstract

Behavioral strategies that depend on sensory information are not immutable; rather they can be shaped by the specific sensory context in which animals develop. This behavioral plasticity depends on the remarkable capacity for the brain to reorganize in response to alterations in the sensory environment, particularly when changes in sensory input occur at an early age. To study this phenomenon, we utilize the short-tailed opossum, a marsupial that has been a valuable animal model to study developmental plasticity due to the extremely immature state of its nervous system at birth. Previous studies in opossums have demonstrated that removal of retinal inputs early in development results in profound alterations to cortical connectivity and functional organization of visual and somatosensory cortex; however, behavioral consequences of this plasticity are not well understood. We trained early blind (EB) and sighted control (SC) opossums to perform a two-alternative forced choice texture discrimination task. Whisker trimming caused an acute deficit in discrimination accuracy for both EB and SC animals indicating that they primarily used a whisker-based strategy to guide choices based on tactile cues – though performance recovered in days, suggesting a shift to the use of other body parts when whiskers were absent. Mystacial whiskers were important for performance in both groups; however, genal whiskers only contributed to performance in EB animals. EB opossums significantly outperformed SC opossums in discrimination accuracy, being more sensitive to textural differences by ~75 μm smaller. Our results support behavioral compensation following early blindness using tactile inputs, especially the whisker system.

## INTRODUCTION

The behavioral strategies that animals use to orient themselves and navigate within complex environments are highly dependent on the sensory context which they inhabit. Over the long timecourse of evolution, different species exhibit sensory adaptations to the ecological niche they occupy. For example, many subterranean mole rats that dwell in burrow systems, with little to no light, are naturally blind, with microphthalmic eyes and no functional vision (Cooper et al., 1993). Instead of vision, they rely heavily on touch and audition, accompanied by a corresponding magnification of sensory representations associated with these senses in cortical and subcortical structures (Mann et al., 1997; Bronchti et al., 2004; Park et al., 2007; Sadka et al., 2004; Kimchi et al., 2004; Kimchi et al., 2005). In fact, what would normally be visual cortex is co-opted by somatosensory and auditory systems. While evolutionary history places some constraints on the sensory-mediated behaviors that animals can engage in, over shorter timescales, behaviors can still be strongly influenced by the sensory context in which an animal develops (Montero, 1997; Arkley et al., 2014). Humans are naturally visual animals (Preuss 2003), however, individuals who experience vision loss, especially in cases of congenital deficits, can become highly effective at using tactile and auditory based strategies along with perceptual learning that manifest as heightened sensitivity to stimuli mediated by the spared senses (Goldreich and Kanics, 2003; Wong et al., 2011; Voss, 2011; for review see Bavelier and Neville, 2002; Kupers and Ptito, 2011; Merabet and Pascual-Leone, 2010; Renier et al., 2014; Ricciardi et al., 2014). Finally, even without the loss of a sensory receptor array, any given individual might rely more heavily on one sensory strategy over another in different settings, depending on the ongoing availability and behavioral relevance of sensory information (Montero, 1997; Lee et al., 2016); for example, touch and hearing may be prioritized over vision upon entering a dark room or at night, when visual information is not readily available.

To appreciate the extent to which early sensory context can shape tactile-mediated behavior, we experimentally altered the relative weights of different sensory inputs in short-tailed opossums through bilateral enucleation at a very early developmental stage (postnatal day 4; P4). P4 in opossums is prior to the onset of spontaneous activity in the retina, and before the formation of retinofugal and thalamocortical pathways (Taylor and Guillery, 1994; Molnar et al., 1998). Previous studies in our laboratory have shown that this manipulation results in a major functional reorganization of visual cortex and alterations in its cortical and subcortical connections (Karlen and Krubitzer, 2009; Karlen et al., 2006; Kahn and Krubitzer, 2002), as well as anatomical and physiological alterations in somatosensory cortex (Dooley and Krubitzer, 2019; Ramamurthy and Krubitzer, 2018; Karlen and Krubitzer, 2009; Karlen et al., 2006; Kahn and Krubitzer, 2002). These changes bear a strong resemblance to cortical organization observed in naturally blind animals (Halley and Krubitzer, 2019). Notably, neurons in primary somatosensory cortex (S1) of early blind opossums are more selective in their responses to single whisker stimuli and showed improved discriminability of whisker identity (Ramamurthy and Krubitzer, 2018). This could support enhanced discrimination of tactile features on a coarse spatial scale, at or above the distance between neighboring whiskers (Diamond et al., 2010). However, it is unknown if early blind opossums are better at sensing and discriminating textures on a very fine spatial scale, and critically, whether this manifests at the behavioral level.

In their natural habitat, short-tailed opossums are semi-arboreal and crepuscular (Eisenberg and Redford, 1989; Carvalho et al., 2011; Macrini, 2004; Jones et al., 2003), preferring low light conditions (Seelke et al., 2014). Under such circumstances, texture becomes especially crucial as a sensory cue, given the paucity of visual cues in dim light. Short-tailed opossums use texture cues to adjust their locomotor kinetics and footfall patterns as behavioral strategies to maintain balance on arboreal substrates (Lammers, 2004; Lammers, 2007; Lammers, 2009; Lammers and Biknevicius, 2004; Lammers et al, 2006; Lammers and Gauntner, 2008). In most small mammals, the facial whiskers are a major channel for gathering sensory information about position, shape and texture of objects in the immediate vicinity of the animal, and are essential for navigation in complicated and irregular settings – especially in the dark (Vincent, 1912; Russell and Pearce, 1970; Diamond et al., 2008; Diamond and Arabzadeh, 2013; Pocock, 1914; Huber, 1930a, 1930b; Lyne, 1959; Ahl, 1986; Sarko et al., 2011; Englund et al., 2020). When opossums are deprived of visual input at an early age, they are forced to become even more heavily reliant on whisker input as the primary means of exploring and navigating their environment (Ramamurthy and Krubitzer, 2018; Ramamurthy et al., 2018; Englund et al., 2020).

In the current study, we examined the role of two sets of facial whiskers – the mystacial whiskers (on the snout) and genal whiskers (on the cheek; Ramamurthy and Krubitzer, 2016) in guiding tactile-based decisions in early blind and sighted opossums, and measured their behavioral discrimination sensitivity to texture cues. To our knowledge, this is the first psychometric investigation of texture discrimination in a marsupial. Moreover, we compare results from sighted animals with those from bilaterally enucleated opossums performing the same task to examine the extent to which developmental history has an impact on texture discrimination sensitivity and the role of facial whiskers in guiding behavior.

## MATERIALS AND METHODS

### Animals

Behavioral experiments were performed in six adult short-tailed opossums (*Monodelphis domestica*; 3 M, 3 F, 4-16 months) obtained from two separate litters. Three of the six animals were bilaterally enucleated at P4 (see below) and were part of a larger study on developmental plasticity of sensory cortex. Where indicated, additional data from pilot experiments in two animals (1M, 1F) are also included. All experimental procedures were approved by UC Davis IACUC and conform to NIH guidelines.

### Bilateral enucleations

Bilateral enucleations were performed at P4 using procedures that have been previously described (Ramamurthy and Krubitzer, 2018). Mothers of experimental litters were lightly anesthetized with Alfaxan (initial dose: 20 mg/kg; maintenance doses: 10-50%; IM) to facilitate enucleation of the pups, which are attached to the mother’s nipples at this developmental stage. Pups were anesthetized by hypothermia. Health of both the mother (respiration rate and body temperature) and the pups (heartbeat, coloration and mobility) was monitored throughout the procedure. An incision was made in the skin covering the eyes, the eyes were removed under microscopic guidance, and the skin was repositioned and resealed using surgical glue. Approximately 50% of each litter was bilaterally enucleated. After complete recovery from anesthesia, mothers along with their attached litters were returned to their home cages.

### Testing apparatus and procedure

Animals were tested in a Y-maze modified to test texture discrimination capabilities using a two-alternative forced choice paradigm (2AFC) (**Figure 1**). A similar approach has previously been used in laboratory rodents (Smith, 1939; Finger and Frommer 1968; Finger et al., 1970; Lipp and Van der Loos, 1991; Hughes, 2007). The Y-maze (**Figure 1A**) was custom-built from plexiglass, and consisted of a corridor (15” long, 12” high) and two arms (12” long, 12” high). A clear plexiglass lid covered the top of the maze. The animal was required to discriminate between a baseline sandpaper (120 grit sandpaper, S+) and a range of sandpapers of different grits (test stimuli, S-). Both the baseline and test stimuli consisted of removable textured panels that could be attached to either arm of the Y-maze, and their respective sides of the corridor. Panels were made of polystyrene and covered with sandpaper of different grit sizes (corresponding to S-stimuli or S+ stimulus). The “goal arm” was defined as the arm which contained the S+ stimulus. On each trial, entry into the “goal arm” was considered a correct choice (hit), while entry into the non-goal arm was considered an incorrect choice (miss). After completion of each trial, the opossum returned to the start zone to begin a new trial. A hit was rewarded with a live cricket. The order of presentation of the different textures was pseudorandomized. 10-12 trials were collected per animal each day. The maze was cleaned with ethanol in between trials to eliminate olfactory cues. If an animal did not consume the reward within one minute following successful trial completion on >50% of hit trials, data from that day was excluded due to low reward-driven motivation for accurately performing the task. All testing was conducted either in dim red light (660 nm – outside the expected peak sensitivity range of cone visual pigments in short-tailed opossums; Hunt et al., 2009) or in the dark, guided by a night vision camera. On a subset of trials, video recordings were captured under infrared lighting using a night vision camera (Seree Night Vision HDV-501, 1080p 60 fps). These were used for analysis of maze exploration and decision latency in opossums.

**Figure 1.**
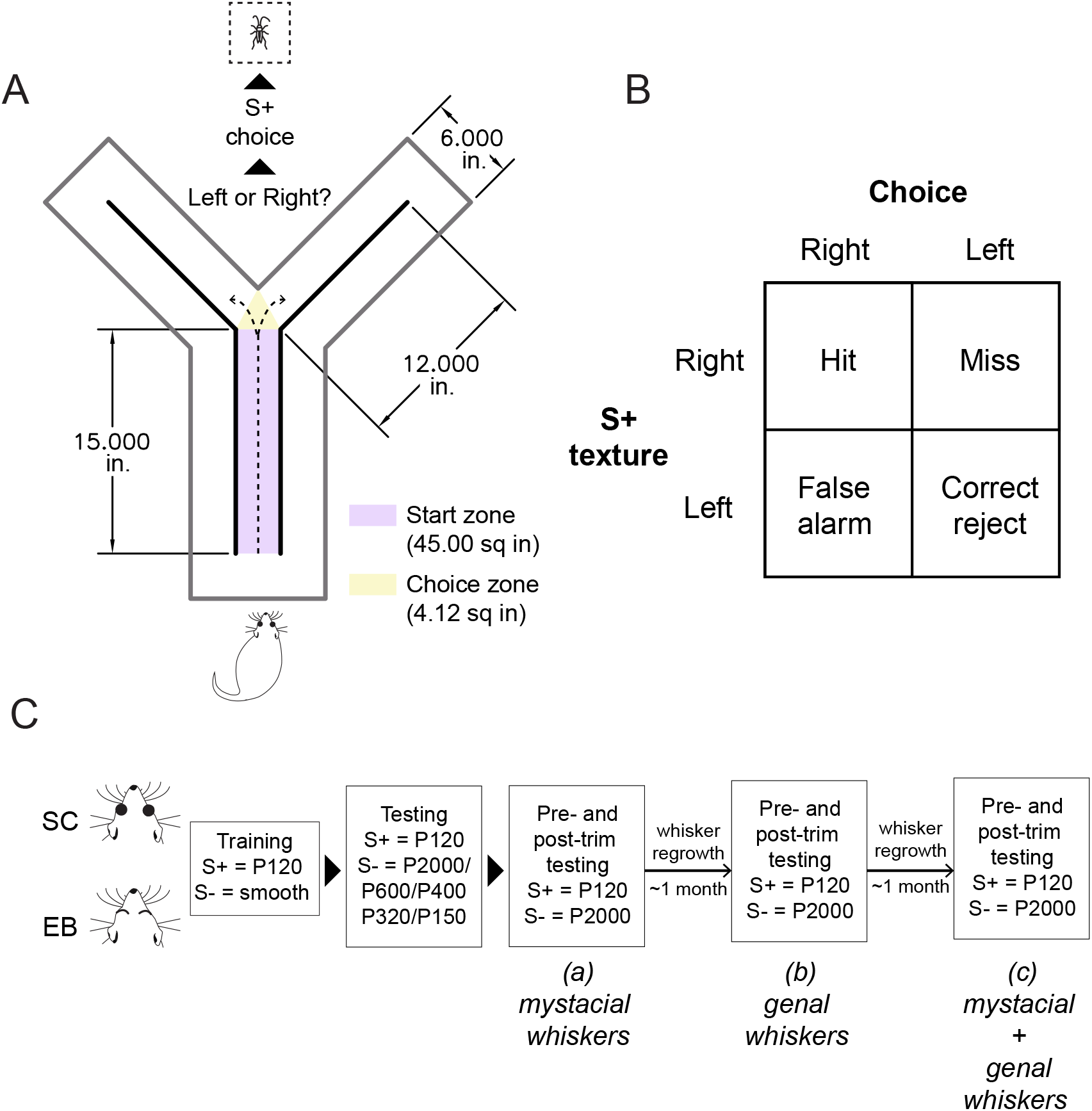
Experimental design **A.** Behavioral testing apparatus. A Y-maze was modified to test texture discrimination capabilities using a two-alternative forced choice paradigm (2AFC). Stimulus panels were attached to the inner walls of the Y-maze (thick black lines). Opossums were required to discriminate between a baseline sandpaper (120 grit sandpaper, S+) and a range of sandpapers of different grits (test stimuli, S-). Choice of the S+ stimulus was rewarded with a live cricket. The start zone and choice zone were designated as shown for the quantification of texture discrimination latency. **B.** Contingency table showing trial types, response types and trial outcomes that result from all possible combinations of trial type and response type. On any given trial, the S+ texture can be presented on the left or right side of the maze (Left/Right trial types) and the animal can respond by choosing either the left or the right arm (Left/Right Choice) in the 2AFC texture discrimination task paradigm. When S+ was presented on the right, choice of the right arm was counted as a hit, while choice of the left arm was a miss. When S+ was presented on the left, choice of the right arm counted as a false alarm, while choice of the left arm was a correct rejection. **C.** Training and testing timeline. Following initial training and testing across all stimulus combinations, the contribution of the facial whiskers to performing this task was determined by trimming (a) mystacial whiskers (b) genal whiskers (c) mystacial and genal whiskers.

### Training stages

Behavior was shaped gradually over a series of training stages described below in sequence.

1. <u>Handling</u>: Handling began immediately following weaning (P56) to facilitate training and testing on the texture discrimination task in adulthood. Handling occurred 2-3 days per week, from weaning until adulthood (see Ramamurthy and Krubitzer, 2018 for developmental timeline). All animals were handled for equal amounts of time (2 minutes) per day. Animals used in pilot experiments did not undergo this initial handling phase.
2. <u>Acclimation to the testing room</u>: Animals were transported to the testing room in their home cages and allowed to acclimate there for around 10 minutes. This phase lasted for 1 day.
3. <u>Acclimation to the testing apparatus</u>: Animals were placed in an open arena in the testing room and then repeatedly moved to and from the testing apparatus (Y-maze) for a total of two minutes per day. This process was repeated 3-5 times a week for 30 days.
4. <u>Stimulus-Reward Pairing</u>: Animals were placed in the open arena, now half covered by the roughest sandpaper (P120 grit). Approach toward and contact with the sandpaper was positively reinforced using a food reward (live cricket), paired with a sound (click). This stage of training lasted for ~2 weeks.
5. <u>2AFC Training</u>: Animals were placed in the start zone (**Figure 1A**) and allowed to navigate the Y-maze. The roughest sandpaper (P120 grit; mean grain size of 125 μm; ISO 6344 industrial standard) was used in the goal arm during this phase. Choice to enter the goal arm was positively reinforced by a click sound, followed by delivery of a cricket as the food reward. This phase lasted until criterion (70% accuracy) was achieved.
6. <u>2AFC Testing</u>: The testing phase began once animals reached criterion performance (70% accuracy). In this phase, the smooth panels were replaced by panels covered with sandpaper of varying roughness (P2000, P600, P400, P320, P150, with mean grain sizes of 10.3 μm, 25.8 μm, 35 μm, 46.2 μm, 100 μm, respectively; ISO 6344 industrial standard). The contrast in textures between the goal arm (120 grit) and the non-goal arm (P120 – P2000) could thus be varied across trials to measure psychometric functions of texture discrimination. Further, data was also acquired on days when either the same texture was presented on both sides, or P120 vs P2000 combination was presented with the panels flipped. This allowed us to control for the possibility of choices being guided by olfactory cues from the sandpaper (Finger et al., 1970). In the flipped panel configuration, the stimulus panels were affixed such that the sandpaper-covered side faced the wall of the Y-maze and the smooth backing of the panel was instead facing inwards, towards the animal. Thus, texture-related cues associated with the stimulus panels were obscured while any odorant cues associated with the sandpaper would be retained. For trials in which either P120 texture was presented in both arms or when flipped panels were used to control for olfactory cues, either the right or the left panel was randomly selected to be the S+ stimulus prior to the start of the session and rewarded accordingly.

### Whisker trimming experiments

Animals were not trained to use any specific body part to perform the texture discrimination. However, the height of the maze was such that it prevented animals from rearing up and actively palpating the stimulus panels with their forepaws. Like mice and rats, short tailed opossums naturally whisk during locomotion and object exploration. The goal of these experiments was to test the extent to which these animals naturally favored the use of their whiskers for texture discrimination to guide behavior.

Following data collection across all tested stimulus combinations, we determined the contribution of the mystacial whiskers (located on the snout) and the genal whiskers (located on the cheek) in performing this task by trimming each set of whiskers separately or at the same time. Three sets of whisker trimming experiments were performed – 1) mystacial whiskers only 2) genal whiskers only 3) both mystacial and genal whiskers. Whiskers were bilaterally trimmed down to approximately 0.5 mm in length (under light isoflurane anesthesia; 1-2% for 3-5 minutes), and the acute effect on behavior was measured for 3-4 days following whisker trimming. Each set of whisker trimming experiments was separated by at least 1 month to allow complete regrowth of all whiskers between experiments. To maintain the stimulus-reward association, some training sessions occurred at <1 month, but these data were not included in the analysis since whisker regrowth could be partial in those cases. 3-5 sessions of data on texture discrimination performance on P120 vs. P2000 combination was acquired pre-trimming, followed by 3-5 sessions for the same stimuli post-trimming. Sham procedures mimicking whisker-trimming (brushing rather than trimming whiskers under 1-2% isoflurane for 3-5 minutes) were included prior to the pre-trimming phase. The first post-trimming behavioral session was conducted no sooner than 18 hours after the whiskers were trimmed.

### Video coding

All video data was manually scored by two independent observers using BORIS (Behavioral Observation Research Interactive) software (version 4.1.4). The parameters scored were: 1) total time spent in the start zone 2) total time spent in the choice zone until the animal left the choice zone and entered one of the arms. The time point of entry into a zone was recorded as the time at which the tip of the snout first entered the zone, while the time point of exit (**Figure 1A**) was recorded as the time at which the whole body of the animal (except the tail) left that zone.

### Data analysis and statistics

All data analyses and statistics were performed in MATLAB version 9.4.0 (MathWorks, Natick, MA) and R version 3.5.2 (R Development Core Team 2018).Texture discrimination performance across different sandpaper combinations was primarily assessed using the metric of accuracy (percent correct trials). Statistical significance of accuracy levels, relative to chance performance was assessed using the binomial exact test. Psychometric functions were plotted for individual animals across different sandpaper combinations. These were used to obtain average psychometric functions, and texture discrimination threshold values across animals in each experimental group and testing condition. Texture discrimination threshold was defined as the smallest difference in sandpaper grit size for which animals showed above chance performance. Mean texture discrimination functions and choice latencies were compared between different experimental groups (SC – sighted controls, and EB – early blind) and experimental conditions (pre-trimming vs. post-trimming; mystacial/genal/both) using two-way ANOVAs. Hit Rate was defined as the proportion of times the animal chose the left arm of the Y-maze, and the S+ stimulus was on the left) (**Figure 1B**). The False Alarm rate is the proportion of times the animal chose the left arm, and the S-stimulus was on the left. The percentage of correct responses was computed as the average of the Hit rate and Correct Rejection rate.

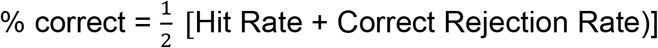

Signal detection theory (Green and Swets, 1966; Carandini and Churchland, 2013) was applied to calculate *d*’ as a measure of discriminability. *d*’ was calculated from the z-score transform of the 2AFC percentage correct (National Research Council Committee on Vision,1985) as follows:

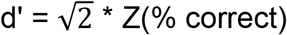

Response bias (b) was calculated using the equation:

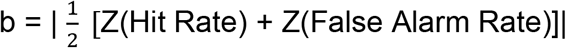

where Z is the inverse cumulative of the normal distribution.

## RESULTS

We trained sighted and early blind short-tailed opossums to perform a two-alternative forced choice (2AFC) texture discrimination task, in which choosing a rougher texture led to a food reward. By varying the difference in roughness between the textures presented for the choice, we generated psychometric functions for texture discrimination in short-tailed opossums and made comparisons of tactile perceptual sensitivity between sighted and early blind experimental groups.

### Sighted and early blind opossums successfully learned to base choices on texture cues to acquire a food reward

In initial training sessions, opossums were presented with only one possible stimulus combination – the S+ stimulus consisting of a P120 sandpaper panel, the choice of which was positively reinforced, and the single S-stimulus consisting of a smooth panel. This was done to benefit task learning by maximizing the textural contrast between S+ and S-stimuli. Animals from both experimental groups achieved criterion-level performance accuracy in choosing the S+ stimulus in ~5 days (**Figure 2A-B**; SC: 5.33 ± 0.27 days; EB: 5.00 ± 0.47 days) after training started (stage 5 of the gradual behavioral shaping process; see *Materials and Methods*). Once the task was learned, both groups of animals reliably spent much more time per unit area exploring in the choice zone of the Y-maze, compared to the start zone (**Figure 2C**; pre-whisker trimming, start zone vs. choice zone – SC: p = 1.26×10^-4^, EB: p = 3.12×10^-4^; paired t-test) before entering into an arm of the maze and terminating the trial, indicating that the animals were engaged in the task, and making choices based on evidence accumulated from sensory cues in the maze. After criterion was attained, on average, performance remained stable for individual animals, and was similar across experimental groups (**Figure 2D**; p = 0.246, 1-way ANOVA; group means: SC: 75.5 ± 2.26%, EB: 77.2 ± 2.19%). To confirm that animals were not guided by possible odorant cues associated with the sandpaper panel, we included a subset of trials where the P120 vs. smooth panel combination was presented with the panels flipped such that texture cues associated with the S+ stimulus were eliminated (see *Materials and Methods*; **Figure 2E**). This caused performance to drop to chance levels in all animals, and was not significantly different from performance when the same texture (P120) was presented on both sides (p = 0.795, 1-way ANOVA; group means, flipped panels – SC: 46.6 ± 3.85%, EB: 51.3 ± 4.12%; group means, both P120 – SC: 45.7 ± 2.75%, EB: 47.8 ± 4.09%). Additionally, we confirmed that no cues associated with the dim red light were guiding choices in sighted animals (**Figure 2F**) by conducting a subset of testing sessions in the dark, under the guidance of only an infrared-enabled camera. Performance was not significantly different under dim red light and dark conditions (p = 0.8917, 1-way ANOVA; group means, red light – SC: 75.2 ± 2.36%, EB: 76.2 ± 1.89%; group means, dark – SC: 75.5 ± 2.34%, EB: 78.1 ± 2.88%). Thus, tactile cues, and not odorant or light cues were responsible for successful task performance in both sighted and blind animals.

**Figure 2.**
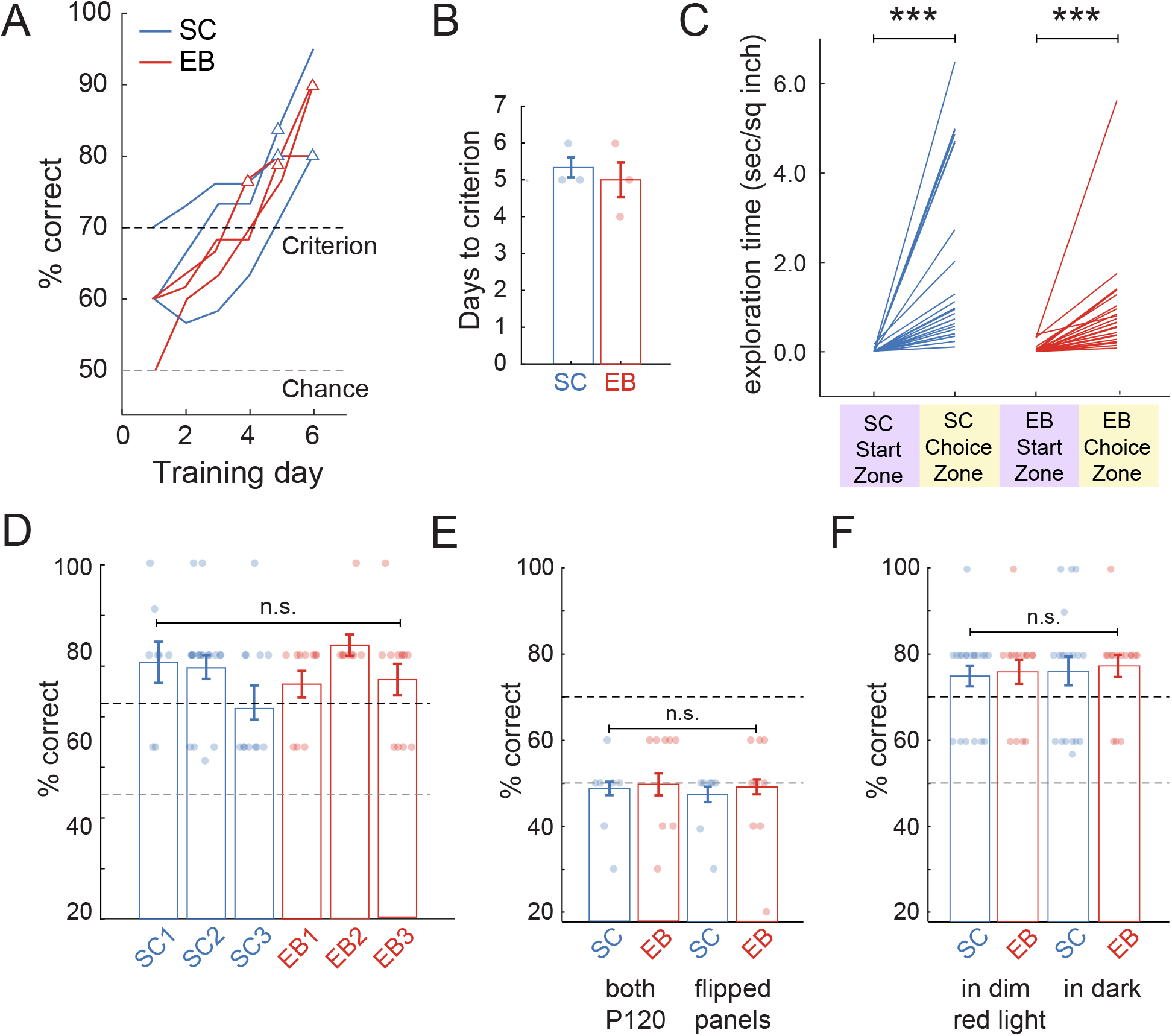
Training results for short-tailed opossums performing the 2AFC texture discrimination task. **A.** Learning curves for four sighted control (SC, blue), and four early blind (EB, red) opossums, for discrimination of the S+ stimulus (P120 sandpaper) from the S-stimulus with maximal difference in texture (smooth panel). Learning curves show moving averages of performance over three days. Animals were considered to reach criterion (triangular markers) after two consecutive training days at ?70%. **B.** Both sighted and early blind opossums attained criterion in ~5 training sessions. **C.** Latency for texture discrimination during performance at criterion. On average, both SC and EB spent significantly more time in the decision zone vs. the initial zone while performing the task, consistent with an active choice of the goal arm rather than selection by chance. **D.** After criterion was reached, P120 vs. smooth discrimination performance was not significantly different between SC and EB groups. **E.** Control for any possible olfactory cues from sandpaper. Presentation of the same texture on both sides (either P120 on both sides or the P120 vs. P2000 combination with flipped panels, smooth side facing in) caused accuracy to drop to chance in both groups. **F.** Control for any possible light cues due to the use of dim red light. Performance in the dark under infrared guidance was not significantly different from performance under dim red light for either group.

### Facial whiskers made a primary contribution in guiding texture-based choices with differential effects of trimming in sighted vs. early blind opossums

We assessed the contribution of facial whiskers to texture discrimination in sighted and early blind short-tailed opossums by trimming the mystacial set of whiskers (located on the snout) and the genal set of whiskers (located on the cheek), either separately or in tandem. Following trimming of all facial whiskers, opossums continued to show similar patterns of exploration in the Y-maze as when whiskers were intact, but they spent significantly greater time per unit area in the choice zone compared to the start zone (**Figure 3A**; post-whisker trimming, start zone vs. choice zone – SC: p = 3.20 x 10^-3^; EB: p = 4.00 x 10^-4;^ paired t-test). For each experimental group, time spent within any particular zone was not significantly different between pre- and post-whisker trimming conditions (pre-vs. post-whisker trimming, start zone – SC: p = 0.896, EB: p = 0.986; choice zone – SC: p = 0.259, EB: p = 0.905; 1-way ANOVA with post-hoc Tukey’s HSD test).

**Figure 3.**
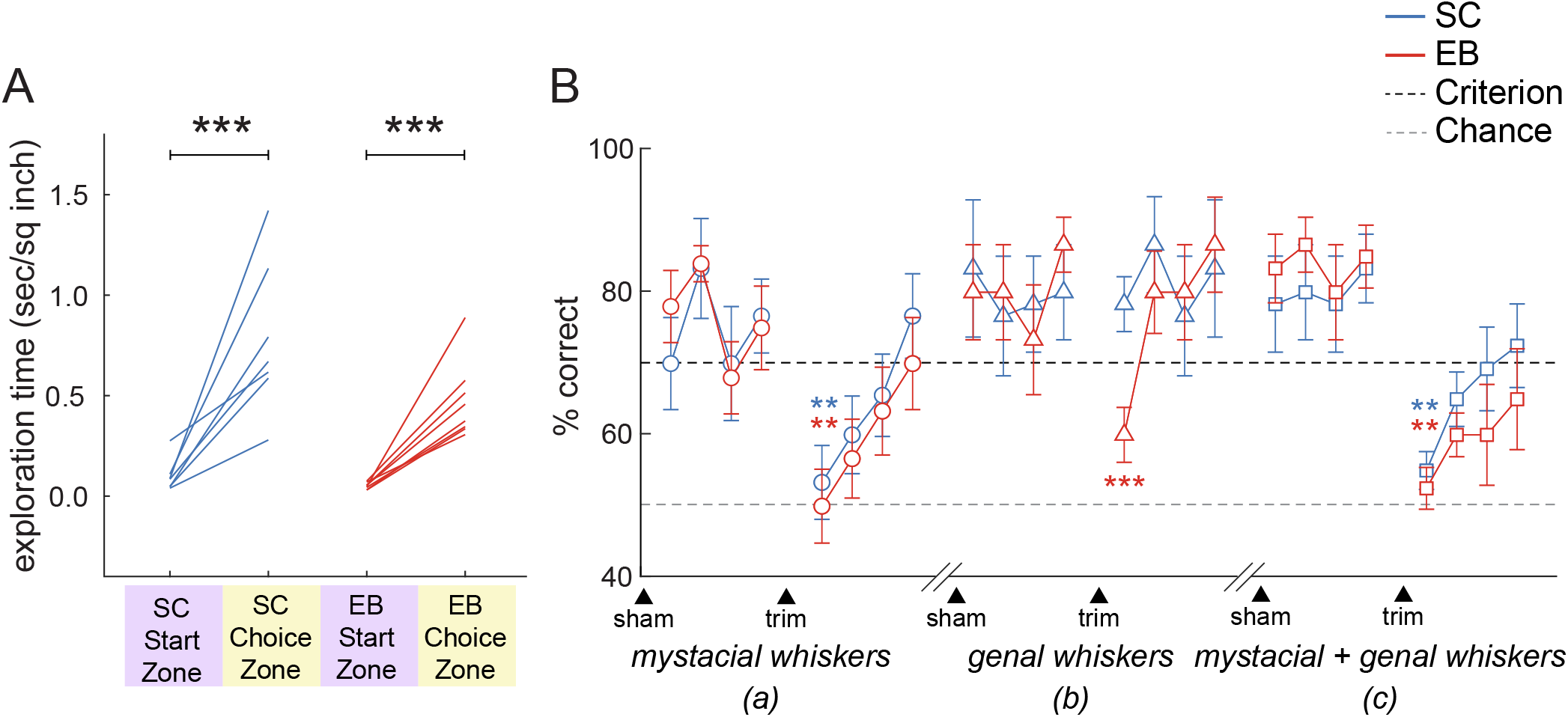
Effects of whisker trimming on texture discrimination task performance. **A.** Latency for texture discrimination after trimming of all facial whiskers. Consistent with their initial training, sighted and early blind animals spent more time in the decision zone vs. the initial zone even after whisker trimming. **B.** Average texture discrimination accuracy across four consecutive testing sessions pre- and post-trimming of the facial whiskers: (a) mystacial whiskers (circles), (b) genal whiskers (triangles), (c) mystacial and genal whiskers (squares). Whisker trimming acutely impaired performance on texture discrimination in both sighted and early blind opossums, which then gradually recovered over the next few sessions. Sham procedures in the pre-trimming phase did not induce a similar drop in performance. Mystacial whisker trimming (either alone or together with genal whisker trimming) diminished texture discrimination accuracy in both experimental groups, while genal whisker trimming affected only the early blind group.

Thus, whisker trimming did not cause opossums to cease being engaged in task performance. Trimming of the mystacial whiskers led to an acute reduction in texture discrimination accuracy on the first day post-trimming and performance dropped to the level of chance in both experimental groups (**Figure 3B**; SC: p = 7.65 x 10^-3^; EB: p = 4.88 x 10^-3^; repeated measures ANOVA with post-hoc Tukey’s HSD test). This was followed by a steady recovery in discrimination accuracy back to criterion levels over the next three days, suggesting that opossums rapidly shifted strategies and relied on other body parts to perform texture discrimination, given that whisker regrowth would be minimal in 1-4 days post-trimming. Notably, trimming of the genal whiskers caused performance to significantly drop on the first day post-trimming only in early blind opossums but not in controls (SC: p = 0.964; EB: p = 4.08 x 10^-7^; repeated measures ANOVA followed by post-hoc Tukey’s HSD test), and remained higher than chance even for the early blind group. By the second day following trimming of the genal whiskers, discrimination performance had fully recovered and styed as such for the next two post-trim days.

Simultaneous trimming of both mystacial and genal whiskers in the third trimming experiment once again caused an acute deficit in whisker discrimination performance on the first day in both groups, similar to what was seen when mystacial whiskers alone were trimmed (SC: p = 1.01 x 10^-2^; EB: p = 7.23 x 10^-3^; repeated measures ANOVA followed by post-hoc Tukey’s HSD test). Importantly, sham procedures mimicking whisker trimming (see *Materials and Methods*), that occurred during the pre-trim phase for each trimming experiment did not cause a dip in performance for either group (**Figure 3B**), indicating that poor performance on the first day after whisker trimming could not be attributed to non-sensory effects such as stress. Thus, when whisker input was present (especially from the mystacial whiskers), it was preferentially used to perform texture-based discriminations. Genal whiskers contributed to performance in early blind animals, but not in sighted animals. When whisker input was removed by trimming, performance recovered for both SC and EB opossums over the time course of a few days, likely through an altered strategy which relied on tactile inputs from other body parts for texture discrimination.

### Early blind opossums showed enhanced texture discrimination accuracy and sensitivity relative to sighted controls

We varied the contrast between the textures of the S+ and S-stimuli in order to generate psychometric functions for texture discrimination in short-tailed opossums. Texture discrimination accuracy dropped with decreases in the difference between two presented textures, quantified as the difference in average grit size between sandpapers (Δ 0-125 μm grit size). For individual animals in both SC and EB experimental groups performance was at chance level for small differences in grit size and increased to criterion level for larger grit size differences (**Figure 4A**). The average texture discrimination curve showed the same generally increasing relationship between textural contrast (Δ sandpaper grit size) and discrimination accuracy for both groups. Importantly, there was a leftward shift in the EB curve relative to the SC curve (**Figure 4B**). There was a significant main effect of grit size difference as well as experimental group on discrimination accuracy (grit size difference: p = 2.19 x10^-10^; experimental group: p = 5.72 x 10^-4^; 2-way ANOVA). On average, early blind opossums could discriminate differences in texture as low as 25 μm (P120 vs. P150; p<0.05, binomial exact test) while sighted controls did no better than chance for differences below Δ 100 μm (P120 vs. P600, p<0.05, binomial exact test). Thus, early blind animals displayed an improved ability to discriminate smaller textural differences, compared to sighted controls under the same conditions.

**Figure 4.**
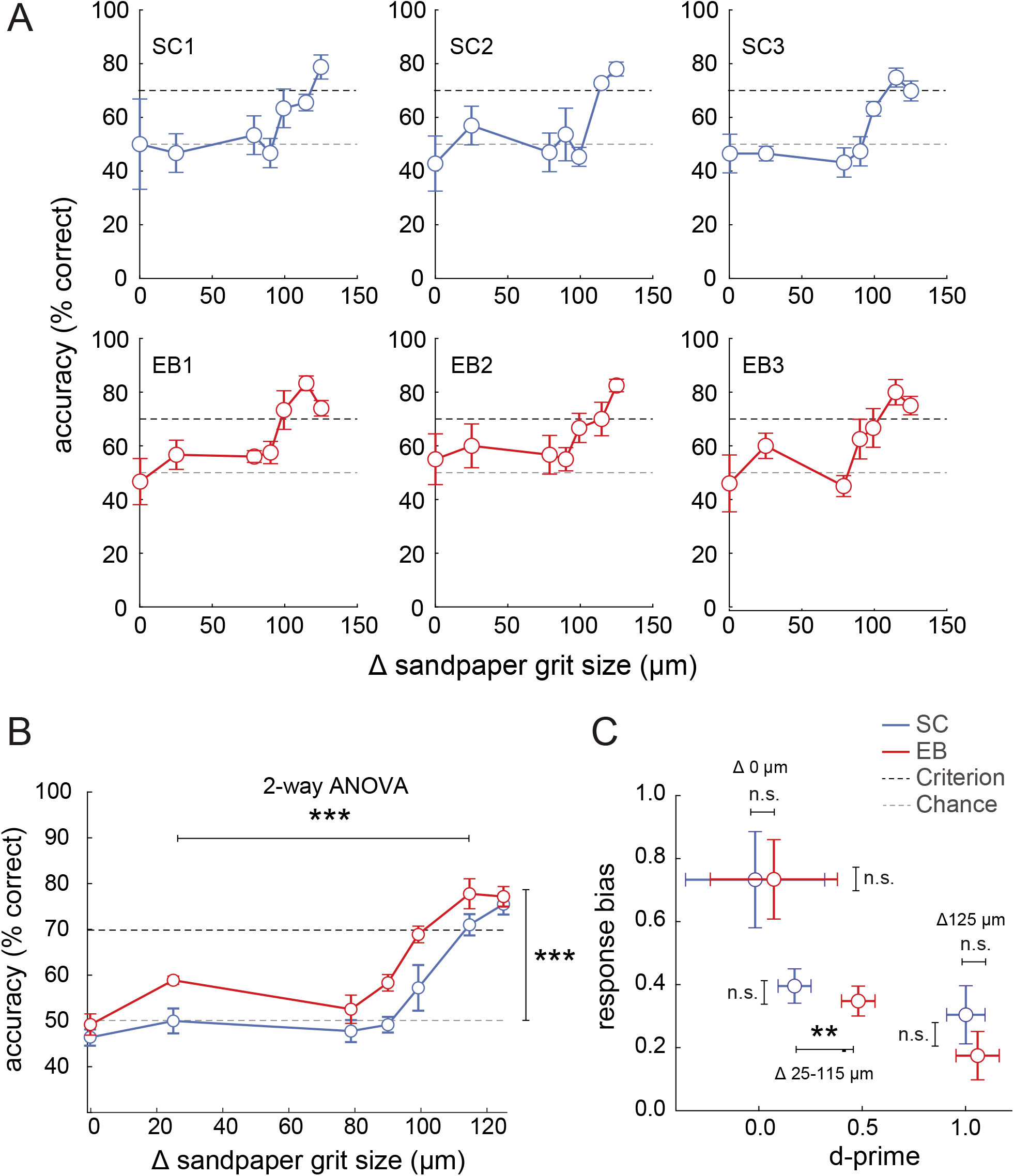
Texture discrimination curves in sighted and early blind opossums **A.** Psychometric functions for three individual sighted control opossums (top row) and three individual early blind opossums (bottom row). All animals have increased discrimination performance as differences in texture increased (Δ sandpaper grit size). **B.** Average psychometric functions for texture discrimination in early blind versus sighted control opossums. Discrimination accuracy increased with texture difference in both SC and EB animals, but the performance curves were shifted to the left for the EB group compared to the SC group, indicating lower texture discrimination thresholds in these animals. Early blind opossums could discriminate differences in texture of as small as 25 μm while sighted controls did no better than chance for < Δ 100 μm differences. **C.** Signal detection theory measures of texture discrimination performance. The scatterplot shows the relationship between mean d-prime and response bias measures for maximal (Δ 125 μm), minimal (Δ 0 μm) and intermediate (Δ 25-115 μm) differences in texture. Only the sensitivity index, d-prime, was significantly different between SC and EB groups, for texture differences of 25-115 μm.

We applied signal detection theory to separate the contributions of sensitivity (*d*’) and bias (*b*) to differences in behavioral performance between SC and EB animals (**Figure 4C**). We computed sensitivity (*d*’) and bias (*b*) when the difference in grit sizes was maximal (125 μm), minimal (0 μm) or in the intermediate range (25-115 μm). Improved texture discrimination accuracy in early blind animals for textural differences in the range of 25-115 μm was accompanied by significant improvements in sensitivity (*d*’) for those stimuli (p = 9.3 x 10^-3^; two sample t-test), but not when the textural difference was maximal (125 μm – initially trained combination, P120 vs. smooth; p = 0.696, two sample t-test) or minimal (0 μm – P120 on both sides, or P120 vs. P2000 flipped panels; p = 0.850, two sample t-test). Response bias (*b*) was inversely related to grit size differences and sensitivity, being high for low Δ grit size stimulus combinations (where sensitivity was low) and low for high Δ grit size stimulus combinations (where sensitivity was high). Response bias did not differ significantly between experimental groups when grit size differences were maximal (p = 0.319; two sample t-test), minimal (p = 0.996; two sample t-test) or in the intermediate range (p = 0.513; two sample t-test). Thus, improved texture discrimination performance for smaller textural differences in early blind opossums can be attributed to improved perceptual sensitivity rather than differences in response bias between the two groups.

## DISCUSSION

### Fine texture discrimination in mammals

Textures are comprised of surface tactile features that can range in scale from coarsely spaced (hundreds of microns to a few millimeters) to finely spaced (<200 microns; Diamond et al., 2010). The discrimination of textures on these different spatial scales involve different neural coding mechanisms, the specifics of which depend on features of any given tactile sensory apparatus – these include the spacing of peripheral receptors, the size of their receptive fields, as well as active touch strategies used (Diamond et al., 2010; Grant et al., 2013). Coarse texture discrimination has been described in a number of mammals that rely on texture for important sensory mediated behaviors using a variety of body parts such as the fingertips in humans (Lamb, 1983; Morley et al., 1983) and non-human primates (squirrel monkey – Hille et al., 2001), the trunk in elephants (Dehnhardt et al., 1997), the forepaws and whiskers in sea otters (Strobel et al., 2018), and the whiskers in harbor seals (Dehnhardt et al., 1998), sea lions (Dehnhardt, 1994; Dehnhardt and Dücker, 1996), manatees (Bachteler and Dehnhardt, 1999; Bauer et al., 2012), and laboratory rodents (Carvell and Simons, 1990). However, fine texture discrimination (Diamond 2010), has only been extensively characterized for the whisker system in laboratory rodents and fingertips in primates (Connor et al., 1990; Connor and Johnson, 1992; Arabzadeh et al., 2005; Hollins and Bensmaia, 2007; von Heimendahl et al., 2007; Wolfe et al., 2008; Diamond 2010; Jadhav and Feldman, 2010; Pacchiarini et al., 2017).

Short-tailed opossums have been shown to use surface texture as a cue to modify their locomotion (Lammers 2009). Here, we found that sighted short-tailed opossums could discriminate between surfaces that differ in roughness by at least ~100 μm (125 μm vs. 25.8 μm mean grit size) while in the dark. Thus, short-tailed opossums are capable of using fine textural differences to guide behavior, as previously reported in rodents (Guic-Robles et al., 1989, Cybulska-Klosowicz and Kossut, 2001, Aggestam and Cahusac, 2007; Bourgeon et al., 2008). However, the smallest textural difference discriminated by sighted opossums in our study is considerably larger than that of rodents. Rats have been reported to discriminate between textures that differ by as low as ~10-20 μm mean grit size (Morita et al., 2011). Mice can discriminate between novel and familiar textures separated by 25 μm in mean grit size (Wu et al., 2013; Wu and Dyck, 2018). It is important to note that the texture difference threshold measured is dependent on the roughness of the base stimulus used, as per Weber’s law – when rats were tested relative to a rougher baseline sandpaper (P150, 100 μm mean grit size), the smallest differences in texture they could discriminate was up to 60 μm larger than when they were tested with a smooth (P1500, 12.6 mm mean grit size) baseline sandpaper (Morita et al., 2011). In our study, test comparisons were made with a relatively rough baseline texture (P120, mean grit size 125 μm); therefore opossums could be capable of discriminating smaller differences in texture than reported here, if a finer grit sandpaper is used as the baseline stimulus.

### Role of the facial whiskers in texture discrimination

Short-tailed opossums have two prominent sets of facial whiskers – the mystacial and genal whiskers, both of which are involved in tactile exploration through active whisking (Grant et al., 2013). We examined the role of these whiskers in the discrimination of textures through whisker-trimming experiments. Both sighted and blind opossums showed diminished performance on the texture discrimination task immediately following trimming of all facial whiskers. This could not be attributed to effects of anesthesia (Lipp and Van der Loos, 1991) or handling procedures associated with trimming because the same procedures minus whisker trimming did not yield a deficit in performance during the pre-trim phase for each set of trimming experiments. Thus, short-tailed opossums used facial whiskers for texture discrimination in this task.

However, trimming of subsets of facial whiskers led to differential effects in EB vs. SC animals. Patterns of performance following mystacial vs. genal whisker trimming indicated that SC animals used only mystacial whiskers for texture discrimination while EB animals used a strategy that integrated sensory information across both mystacial and genal whiskers. This suggests that even under normal circumstances mystacial whiskers are involved in fine texture discrimination but genal whiskers may perform different sensory functions than mystacial whiskers, as has been reported for other groups of whiskers in other mammals (for example, the whisker trident in rats; Thé et al., 2013; Chorev et al., 2016). The differential contribution of mystacial and genal whiskers to performance could be due to differences in active sampling strategies engaged during whisking and locomotion in blind vs. sighted animals (Arkley et al., 2014) developed to compensate for the lack of vision. With the loss of a major sensory system (vision) genal whiskers appear to be recruited for making fine texture discriminations, indicating that the strategy employed for adaptive sensory mediated behaviors is highly flexible and dependent on available inputs from the different sensory systems.

There is evidence for behavioral flexibility in sighted animals as well. In all trimming experiments for both experimental groups, texture discrimination performance recovered over the next three days when the trimmed whiskers had still not grown back. Given that we verified that olfactory and visual cues were not used to perform the discrimination task (**Figure 2E-F**), it appears that both SC and EB animals utilized strategies mediated by other body parts for making tactile discriminations. This could have involved the use of microvibrissae (Kerekes et al., 2017; Kuruppath et al., 2014; Morita et al., 2011), skin on the snout (Kerekes et al., 2017; Morita et al., 2011) or possibly even skin or fur on the trunk (Kerekes et al., 2017). Quick recovery of performance in the post-trim phase has also been reported in rats allowed to freely run while discriminating fine tactile patterns (Kerekes et al., 2017). Notably, once whiskers had fully grown back, opossums returned to using a whisker-based strategy during the task – as was evident from the reduction in performance seen once again in the post-trim phase following the last trimming experiment, when both sets of facial whiskers were trimmed. In the absence of visual information either temporarily (in the case of SC animals) or over the course of development (in the case of EB animals), opossums favored a whisker-based strategy to discriminate between textures. Thus, while opossums were predisposed to using their whiskers for texture discrimination, they could flexibly recruit alternative strategies based on available sensory inputs, over the course of their lifetime. Such behavioral flexibility, which is likely cortically mediated, is a common feature in mammals (Krubitzer and Prescott, 2018).

### Enhanced tactile behavioral sensitivity following vision loss

In the current study, we demonstrate that texture discrimination performance in short-tailed opossums can be altered by developmental history – for the same sandpaper combinations, early blind opossums discriminated differences in textures by as much as ~75 μm lower than the smallest discrimination made by sighted controls (EB: 25 μm vs. SC: 100 μm). Several studies have reported enhanced tactile perception in early blind human subjects (Walker and Moylan, 1993; Goldreich & Kanics, 2003; Goldreich & Kanics, 2006; Legge et al., 2008; Alary et al., 2009; Wong et al., 2011), although this was not demonstrated in all studies (Heller 1989; Alary et al., 2009; Gurtubay-Antolin and Rodríguez-Fornells, 2017). Studies comparing different touch-based tasks revealed enhancement in sensory performance for some tasks but not others (Alary et al., 2009; Gurtubay-Antolin and Rodríguez-Fornells, 2017). Further, other studies have shown that even for tasks in which early blind subjects showed superior tactile performance, this was seen for some body parts, but not others (Wong et al., 2011). Specifically, tactile spatial acuity was greater for the preferred reading finger of Braille readers compared to a nonpreferred fingers or other body parts, and was correlated with the level of use. These findings support the contribution of use-dependent mechanisms to the development of heightened performance via the spared senses.

Thus, whether or not a specific aspect of tactile performance is enhanced in blind individuals could depend on the behavioral strategies that they used to engage with their environment (Arkley et al 2014; Schinazi et al., 2016), among other factors. This can be especially impactful over the course of development when experience can have a major influence on plasticity in the nervous system. Given that even sighted opossums were found to use a primarily mystacial whisker-dependent strategy to solve the texture discrimination task in the dark, it follows that increased reliance on the whiskers to perform this function in the absence of vision from an early age could result in use-dependent plasticity. Our study shows that opossums that are blind from a very early stage in development are capable of enhanced discrimination of finely scaled textures using a primarily whisker-dependent strategy. These findings add to our previous results from recordings in primary somatosensory cortex of early blind short-tailed opossums which provided evidence of enhanced neural discrimination at a coarser spatial scale, for tactile stimuli applied to different whiskers on the face (Ramamurthy and Krubitzer, 2018). In that study we found that S1 neurons in EB animals were more selective in their responses to the deflection of individual whiskers, especially along the rostrocaudal axis of the snout, in alignment with the primary axis of natural whisker motion. Together, these studies provide support for enhancement of the representation of tactile information across multiple spatial scales in short-tailed opossums following the loss of vision early in development.

## ACKNOWLEDGMENTS

We thank Mackenzie Englund, Andrew Halley, Carlos Pineda and Chris Bresee for comments on a draft of this manuscript, and Dr. Cindy Clayton for animal care.

## AUTHOR CONTRIBUTIONS

Conceptualization: D.L.R, L.A.K; Methodology: D.L.R, H.K.D, L.A.K; Formal analysis: D.L.R, H.K.D; Investigation: D.L.R, H.K.D; Writing - original draft: D.L.R; Writing - review & editing: D.L.R, H.K.D, L.A.K; Visualization: D.L.R.; Supervision: L.A.K; Project administration: D.L.R, L.A.K; Funding acquisition: D.L.R, L.A.K.

## REFERENCES

Aggestam, F. and Cahusac, P. M. (2007). Behavioural lateralization of tactile performance in the rat. Physiol Behav 91, 335–9.

Ahl, A. (1986). The role of vibrissae in behavior: a status review. Veterinary research communications 10, 245–268.

Alary, F., Duquette, M., Goldstein, R., Elaine Chapman, C., Voss, P., La Buissonnière-Ariza, V. and Lepore, F. (2009). Tactile acuity in the blind: A closer look reveals superiority over the sighted in some but not all cutaneous tasks. Neuropsychologia 47, 2037–2043.

Arabzadeh, E., Zorzin, E. and Diamond, M. E. (2005). Neuronal Encoding of Texture in the Whisker Sensory Pathway. PLOS Biology 3, e17.

Arkley, K., Grant, R. A., Mitchinson, B. and Prescott, T. J. (2014). Strategy change in vibrissal active sensing during rat locomotion. Curr Biol 24, 1507–12.

Bachteler D, Dehnhardt G (1999). Active touch performance in the Antillean manatee: evidence for a functional differentiation of facial tactile hairs. Zoology 102, 61–69

Bauer, G. B., Gaspard III, J. C., Colbert, D. E., Leach, J. B., Stamper, S. A., Mann, D. and Reep, R. (2012). Tactile discrimination of textures by Florida manatees (Trichechus manatus latirostris). Marine Mammal Science 28, E456–E471.

Bavelier D., Neville H.J. (2002). Cross-modal plasticity: where and how? Nature reviews Neuroscience 3, 443–452.

Bourgeon, S., Xerri, C. and Coq, J.-O. (2004). Abilities in tactile discrimination of textures in adult rats exposed to enriched or impoverished environments. Behavioural brain research 153, 217–231.

Bronchti G., Heil P., Sadka R., Hess A., Scheich H., Wollberg Z. (2002). Auditory activation of “visual” cortical areas in the blind mole rat (Spalax ehrenbergi). The European journal of neuroscience 16, 311–329.

Carandini, M. and Churchland, A. K. (2013). Probing perceptual decisions in rodents. Nat Neurosci 16, 824–31.

Carvalho, B. d. A., Oliveira, L. F. B., Langguth, A., Freygang, C. C., Ferraz, R. S. and Mattevi, M. S. (2011). Phylogenetic relationships and phylogeographic patterns in Monodelphis (Didelphimorphia: Didelphidae). Journal of Mammalogy 92, 121–133.

Carvell, G. E. and Simons, D. J. (1990). Biometric analyses of vibrissal tactile discrimination in the rat. J Neurosci 10, 2638–48.

Chorev, E., Preston-Ferrer, P. and Brecht, M. (2016). Representation of egomotion in rat’s trident and E-row whisker cortices. Nat Neurosci 19, 1367–73.

Connor, C. E., Hsiao, S. S., Phillips, J. R. and Johnson, K. O. (1990). Tactile roughness: neural codes that account for psychophysical magnitude estimates. The Journal of Neuroscience 10, 3823.

Connor, C. E. and Johnson, K. O. (1992). Neural coding of tactile texture: comparison of spatial and temporal mechanisms for roughness perception. The Journal of Neuroscience 12, 3414.

Cooper, H. M., Herbin, M. and Nevo, E. (1993). Visual system of a naturally microphthalmic mammal: the blind mole rat, Spalax ehrenbergi. J Comp Neurol 328, 313–50.

Cybulska-Klosowicz, A. and Kossut, M. (2001). Mice can learn roughness discrimination with vibrissae in a jump stand apparatus. Acta Neurobiol Exp (Wars) 61, 73–6.

Dehnhardt, G., Friese, C. and Sachser, N. (1997). Sensitivity of the trunk of Asian elephants for texture differences of actively touched objects. Z. Saugetierkd. 62, 37–39.

Dehnhardt, G; Mauck, B and Bleckmann, H (1998). Seal whiskers detect water movements. Nature 394, 235–236.

Dehnhardt, G. (1994). Tactile size discrimination by a California sea lion (Zalophus californianus) using its mystacial vibrissae. Journal of Comparative Physiology A 175, 791–800.

Dehnhardt, G. and Dücker, G. (1996). Tactual discrimination of size and shape by a California sea lion (Zalophus californianus). Animal Learning & Behavior 24, 366–374.

Diamond, M. E. (2010). Texture sensation through the fingertips and the whiskers. Curr Opin Neurobiol 20, 319–27.

Diamond, M. E. and Arabzadeh, E. (2013). Whisker sensory system – From receptor to decision. Progress in Neurobiology 103, 28–40.

Diamond, M. E., von Heimendahl, M., Knutsen, P. M., Kleinfeld, D. and Ahissar, E. (2008). ‘Where’ and ‘what’ in the whisker sensorimotor system. Nature Reviews Neuroscience 9, 601–612.

Dooley, J. C. and Krubitzer, L. A. (2019). Alterations in cortical and thalamic connections of somatosensory cortex following early loss of vision. Journal of Comparative Neurology 527, 1675–1688.

Englund, M., Faridjoo, S., Iyer, C. and Krubitzer, L. (2020). Performance and behavioral flexibility on a complex motor task depend on available sensory inputs in early blind and sighted short-tailed opossums. bioRxiv, 2020.05.12.091108.

Eisenberg, J. F. and Redford, K. H. (1989). Mammals of The Neotropics, Volume 3: The Central Neotropics: Ecuador, Peru, Bolivia, Brazil. In Mammals of The Neotropics, Volume 3: The Central Neotropics: Ecuador, Peru, Bolivia, Brazil.

Finger, S. and Frommer, G. P. (1968). Effects of cortical lesions on tactile discriminations graded in difficulty. Life Sciences 7, 897–904.

Finger, S., Frommer, G. P., Carmon, A. and Inbal, R. (1970). Roughness discrimination with sandpaper surfaces: An olfactory confounding. Psychonomic Science 18, 165–166.

Goldreich D., Kanics I.M. (2003). Tactile acuity is enhanced in blindness. J Neurosci 23:3439–3445.

Grant, R. A., Haidarliu, S., Kennerley, N. J. and Prescott, T. J. (2013). The evolution of active vibrissal sensing in mammals: evidence from vibrissal musculature and function in the marsupial opossum *Monodelphis domestica*. The Journal of Experimental Biology 216, 3483–3494.

Green, D. M. and Swets, J. A. (1966). Signal detection theory and psychophysics. Oxford, England: John Wiley.

Guic-Robles, E., Valdivieso, C. and Guajardo, G. (1989). Rats can learn a roughness discrimination using only their vibrissal system. Behav Brain Res 31, 285–9.

Gurtubay-Antolin, A. and Rodríguez-Fornells, A. (2017). Neurophysiological evidence for enhanced tactile acuity in early blindness in some but not all haptic tasks. NeuroImage 162, 23–31.

Halley, A. C. and Krubitzer, L. (2019). Not all cortical expansions are the same: the coevolution of the neocortex and the dorsal thalamus in mammals. Curr Opin Neurobiol 56, 78–86.

Heller, M. A. (1989). Texture perception in sighted and blind observers. Perception & Psychophysics 45, 49–54.

Hille, C. B.-C. G. D. G. D. P. (2001). Haptic discrimination of size and texture in squirrel monkeys ( Saimiri sciureus). Somatosensory & Motor Research 18, 50–61.

Hollins, M. and Bensmaïa, S. J. (2007). The coding of roughness. Canadian Journal of Experimental Psychology/Revue canadienne de psychologie expérimentale 61, 184–195.

Huber E. (1930a). Evolution of facial musculature and cutaneous field of trigeminus. Part I. Q Rev Biol 5, 133–188.

Huber E. (1930b). Evolution of facial musculature and cutaneous field of trigeminus. Part II. Q Rev Biol 5, 389–437.

Hughes, R. N. (2007). Rats’ responsiveness to tactile changes encountered in the dark, and the role of mystacial vibrissae. Behavioural Brain Research 179, 273–280.

Hunt, D. M., Chan, J., Carvalho, L. S., Hokoc, J. N., Ferguson, M. C., Arrese, C. A. and Beazley, L. D. (2009). Cone visual pigments in two species of South American marsupials. Gene 433, 50–5.

Jadhav, S. P. and Feldman, D. E. (2010). Texture coding in the whisker system. Curr Opin Neurobiol 20, 313–8.

Jones, M., Archer, M. and Dickman, C. (2003). Predators with Pouches: The Biology of Carnivorous Marsupials: CSIRO Publishing.

Kahn, D. M. and Krubitzer, L. (2002). Massive cross-modal cortical plasticity and the emergence of a new cortical area in developmentally blind mammals. Proceedings of the National Academy of Sciences 99, 11429–11434.

Karlen SJ, Kahn DM, Krubitzer L. (2006). Early blindness results in abnormal corticocortical and thalamocortical connections. Neuroscience. 142, 843–858.

Karlen, S. J. and Krubitzer, L. (2009). Effects of bilateral enucleation on the size of visual and nonvisual areas of the brain. Cerebral cortex (New York, N.Y.: 1991) 19, 1360–1371.

Kerekes, P., Daret, A., Shulz, D. E. and Ego-Stengel, V. (2017). Bilateral Discrimination of Tactile Patterns without Whisking in Freely Running Rats. The Journal of Neuroscience 37, 7567.

Kimchi, T. and Terkel, J. (2004). Comparison of the role of somatosensory stimuli in maze learning in a blind subterranean rodent and a sighted surface-dwelling rodent. Behavioural brain research 153, 389–395.

Kimchi, T., Reshef, M. and Terkel, J. (2005). Evidence for the use of reflected self-generated seismic waves for spatial orientation in a blind subterranean mammal. Journal of Experimental Biology 208, 647–659.

Krubitzer, L. A. and Prescott, T. J. (2018). The Combinatorial Creature: Cortical Phenotypes within and across Lifetimes. Trends Neurosci 41, 744–762.

Kupers, R. and Ptito, M. (2011). Insights from darkness: what the study of blindness has taught us about brain structure and function. In Progress in brain research, vol. 192, pp. 17–31: Elsevier.

Kuruppath, P., Gugig, E. and Azouz, R. (2014). Microvibrissae-based texture discrimination. J Neurosci 34, 5115–20.

Lamb, G. D. (1983). Tactile discrimination of textured surfaces: psychophysical performance measurements in humans. J Physiol 338, 551–65.

Lammers, A. R. (2004). The biodynamics of arboreal locomotion in the gray short-tailed opossum (Monodelphis domestica).

Lammers, A. R. (2007). Locomotor kinetics on sloped arboreal and terrestrial substrates in a small quadrupedal mammal. Zoology (Jena) 110, 93–103.

Lammers, A. R. (2009). The effects of substrate texture on the mechanics of quadrupedal arboreal locomotion in the gray short-tailed opossum (Monodelphis domestica). J Exp Zool A Ecol Genet Physiol 311, 813–23.

Lammers, A. R. and Biknevicius, A. R. (2004). The biodynamics of arboreal locomotion: the effects of substrate diameter on locomotor kinetics in the gray short-tailed opossum (Monodelphis domestica). J Exp Biol 207, 4325–36.

Lammers, A. R., Earls, K. D. and Biknevicius, A. R. (2006). Locomotor kinetics and kinematics on inclines and declines in the gray short-tailed opossum Monodelphis domestica. J Exp Biol 209, 4154–66.

Lammers, A. R. and Gauntner, T. (2008). Mechanics of torque generation during quadrupedal arboreal locomotion. Journal of Biomechanics 41, 2388–2395.

Lee, C. C. Y., Diamond, M. E. and Arabzadeh, E. (2016). Sensory Prioritization in Rats: Behavioral Performance and Neuronal Correlates. The Journal of Neuroscience 36, 3243–3253.

Legge, G. E., Madison, C., Vaughn, B. N., Cheong, A. M. Y. and Miller, J. C. (2008). Retention of high tactile acuity throughout the life span in blindness. Perception & Psychophysics 70, 1471–1488.

Lipp, H. P. and Van der Loos, H. (1991). A computer-controlled Y-maze for the analysis of vibrissotactile discrimination learning in mice. Behav Brain Res 45, 135–45.

Lyne AG. (1959). The systematic and adaptive significance of the vibrissae in the Marsupialia. Proc Zool Soc Lond 133, 79–133.

Macrini, T. E. (2004). Monodelphis domestica. Mammalian Species, 1–8.

Mann, M. D., Rehkämper, G., Reinke, H., Frahm, H. D., Necker, R. and Nevo, E. (1997). Size of somatosensory cortex and of somatosensory thalamic nuclei of the naturally blind mole rat, Spalax ehrenbergi. J Hirnforsch 38, 47–59.

Merabet, L. B. and Pascual-Leone, A. (2010). Neural reorganization following sensory loss: the opportunity of change. Nature Reviews Neuroscience 11, 44–52.

Morita, T., Kang, H., Wolfe, J., Jadhav, S. P. and Feldman, D. E. (2011). Psychometric Curve and Behavioral Strategies for Whisker-Based Texture Discrimination in Rats. PLOS ONE 6, e20437.

Morley, J. W., Goodwin, A. W. and Darian-Smith, I. (1983). Tactile discrimination of gratings. Experimental Brain Research 49, 291–299.

Molnar, Z., Knott G. W., Blakemore C., Saunders, N. R. (1998). Development of thalamocortical projections in the South American gray short-tailed opossum (Monodelphis domestica). J Comp Neurol 398, 491–514.

Montero, V. M. (1997). c-fos induction in sensory pathways of rats exploring a novel complex environment: shifts of active thalamic reticular sectors by predominant sensory cues. Neuroscience 76, 1069–81.

National Research Council Committee on Vision. (1985). Emergent techniques for the assessment of visual performance. Washington, DC: National Academy Press.

Pacchiarini, N., Fox, K. and Honey, R. C. (2017). Perceptual learning with tactile stimuli in rodents: Shaping the somatosensory system. Learning & Behavior 45, 107–114.

Park, T. J., Catania, K. C., Samaan, D. and Comer, C. M. (2007). Adaptive Neural Organization of Naked Mole-Rat Somatosensation (and Those Similarly Challenged). In Subterranean Rodents: News from Underground, eds. S. Begall H. Burda and C. E. Schleich), pp. 175–193. Berlin, Heidelberg: Springer Berlin Heidelberg.

Preuss, T. M. (2003). 10 Specializations of the Human Visual System: The Monkey Model Meets Human Reality. The primate visual system, 231.

Pocock, R. (1914). 48. On the Facial Vibrissæ of Mammalia. Journal of Zoology 84, 889–912.

R: A language and environment for statistical computing. R Foundation for Statistical Computing, Vienna, Austria. http://www.R-project.org/.

Ramamurthy, D. L. and Krubitzer, L. A. (2016). The evolution of whisker-mediated somatosensation in mammals: Sensory processing in barrelless S1 cortex of a marsupial, Monodelphis domestica. J Comp Neurol 524, 3587–3613.

Ramamurthy, D. L. and Krubitzer, L. A. (2018). Neural Coding of Whisker-Mediated Touch in Primary Somatosensory Cortex Is Altered Following Early Blindness. The Journal of Neuroscience 38, 6172–6189.

R Core Team. (2018).

Renier L., De Volder A.G., Rauschecker J.P. (2014) Cortical plasticity and preserved function in early blindness. Neurosci Biobehav Rev 41:53–63.

Ricciardi E., Handjaras G., Pietrini P. (2014) The blind brain: how (lack of) vision shapes the morphological and functional architecture of the human brain. Exp Biol Med (Maywood) 239:1414–1420.

Russell, E. M. and Pearce, G. A. (1971). Exploration of Novel Objects by Marsupials. Behaviour 40, 312–322.

Sadka R.S., Wollberg Z. (2004). Response properties of auditory activated cells in the occipital cortex of the blind mole rat: an electrophysiological study. J Comp Physiol A Neuroethol Sens Neural Behav Physiol 190, 403–413.

Sarko, D. K., Ghose, D. and Wallace, M. T. (2013). Convergent approaches toward the study of multisensory perception. Front Syst Neurosci 7, 81.

Schinazi, V. R., Thrash, T. and Chebat, D.-R. (2016). Spatial navigation by congenitally blind individuals. Wiley interdisciplinary reviews. Cognitive science 7, 37–58.

Seelke, A. M., Dooley, J. C. and Krubitzer, L. A. (2014). Photic preference of the short-tailed opossum (Monodelphis domestica). Neuroscience 269, 273–80.

Smith, D. E. (1939). Cerebral localization in somaesthetic discrimination in the rat. Journal of Comparative Psychology 28, 161–188.

Strobel, S. M., Sills, J. M., Tinker, M. T. and Reichmuth, C. J. (2018). Active touch in sea otters: in-air and underwater texture discrimination thresholds and behavioral strategies for paws and vibrissae. The Journal of Experimental Biology 221, jeb181347.

Taylor, J. S. H. and Guillery, R. W. (1994). Early development of the optic chiasm in the gray short-tailed opossum, Monodelphis domestica. Journal of Comparative Neurology 350, 109–121.

Thé, L., Wallace, M. L., Chen, C. H., Chorev, E. and Brecht, M. (2013). Structure, function, and cortical representation of the rat submandibular whisker trident. J Neurosci 33, 4815–24.

Vincent, S. B. (1912). The function of the vibrissae in the behavior of the white rat. Animal Behavior Monographs 1, 5, 84–84.

von Heimendahl, M., Itskov, P. M., Arabzadeh, E. and Diamond, M. E. (2007). Neuronal Activity in Rat Barrel Cortex Underlying Texture Discrimination. PLOS Biology 5, e305.

Voss P. (2011). Superior Tactile Abilities in the Blind: Is Blindness Required? The Journal of Neuroscience 31, 11745–11747.

Walker, P. and Moylan, K. (1994). The enhanced representation of surface texture consequent on the loss of sight. Neuropsychologia 32, 289–297.

Wolfe, J., Hill, D. N., Pahlavan, S., Drew, P. J., Kleinfeld, D. and Feldman, D. E. (2008). Texture Coding in the Rat Whisker System: Slip-Stick Versus Differential Resonance. PLOS Biology 6, e215.

Wong M., Gnanakumaran V., Goldreich D. (2011) Tactile spatial acuity enhancement in blindness: evidence for experience-dependent mechanisms. J Neurosci 31, 7028–7037

Wu, H.-P. P., and Dyck, R. H. (2018). Signaling by Synaptic Zinc is Required for Whisker-Mediated, Fine Texture Discrimination. Neuroscience 369, 242–247.

Wu, H.-P. P., Ioffe, J. C., Iverson, M. M., Boon, J. M. and Dyck, R. H. (2013). Novel, whisker-dependent texture discrimination task for mice. Behavioural brain research 237, 238–242.

